# An essential role for the Hv1 voltage-gated proton channel in *Pseudomonas aeruginosa* corneal infection

**DOI:** 10.1101/2024.07.15.603631

**Authors:** Priscila Rodriguez, Serena Abbondante, Michaela Marshall, Jolynn Tran Chau, Jessica Abdelmeseh, Jamison L. Nourse, Francesco Tombola, Eric Pearlman

## Abstract

Assembly of NADPH oxidase 2 (NOX2) proteins in neutrophils plays an essential role in controlling microbial infections by producing high levels of reactive oxygen species (ROS). We reported that neutrophils and NOX2 are required to control *P. aeruginosa* in a clinically relevant murine model of blinding corneal infection. Given the published role for the voltage-gated proton channel Hv1 in sustaining NOX2 production, we examined the role of Hv1 in *P. aeruginosa* keratitis. *Hvcn1^−/−^* mice exhibited an impaired ability to kill bacteria that was associated with reduced neutrophil recruitment to infected corneas. Unlike earlier reports, we found that *Hvcn1^−/−^* neutrophils produce more rather than less ROS compared with control neutrophils infected with *P. aeruginosa* or stimulated with PMA or zymosan. Collectively, we demonstrate that Hv1 has an important role in control of bacterial growth by neutrophils in bacterial infection beyond the regulation of ROS production.

## Introduction

Microbial infections of the cornea are among the leading causes of preventable blindness globally [1]. Although multiple organisms can infect the cornea, *Pseudomonas aeruginosa* is a major cause of corneal infections in the USA and worldwide. Poor contact lens hygiene is an important risk factor in industrialized countries, whereas ocular trauma is the major predisposing cause in developing countries [1]. These infections result in corneal opacity, inflammation, and intense pain, and can lead to permanent blindness if left untreated. While antibiotics are the first line of defense, there are increasing reports of antibiotic resistance in clinical isolates of *P. aeruginosa*, including a 2023 outbreak of drug-resistant *P. aeruginosa* keratitis due to a contaminated artificial tears product that was imported to the USA. There were 68 patients from 16 states who were infected with this virulent strain, resulting in 3 deaths and 8 enucleations [2, 3]. Although these eyedrops have been withdrawn from use, this outbreak illustrates the importance of developing antimicrobial and anti-inflammatory therapies as corneal opacification is also due to cellular infiltration and inflammation.

Our collaborative studies at the Aravind Eye Hospital in Tamil Nadu, India identified neutrophils as the major cell type (>90%) infiltrating corneal ulcers caused by *P. aeruginosa, Streptococcus pneumoniae* or pathogenic fungi [4, 5]. We also demonstrated that neutrophils are the predominant cell type in infected corneas in a murine models of *P. aeruginosa* keratitis, and that depletion of neutrophils results in unimpaired bacterial growth and corneal ulceration [6, 7].

Neutrophils utilize both oxidative and non-oxidative effectors to kill bacteria, including iron and zinc binding proteins that compete with bacterial siderophores and transporters, potent phagocytic activity and release of extracellular traps containing DNA and microbicidal histones [8, 9]. However, reactive oxygen species (ROS) play an outsized role in bacterial killing, especially following phagocytosis and formation of phagolysosomes.

Neutrophil ROS production in phagolysosomes is primarily generated by the NADPH oxidase protein complex 2 (NOX2) that includes the gp91^phox^ and p22^phox^ proteins on plasma and phagosome membranes, which together with the Rac GTPase comprise the flavocytochrome b558 (cytb558) subunit. The regulatory p40^phox^, p47^phox^ and p67^phox^ proteins in the cytoplasm translocate to the membrane following phosphorylation of p47^phox^ and associate with cytb558 to form NOX2 (reviewed in [10]). Once assembled, NOX2 generates ROS superoxide (O_2_^−^) by accepting electrons from cytoplasmic NADPH and donating them to molecular oxygen, leading to membrane depolarization and accumulation of protons in the cytoplasm, which in turn leads to acidification of the cytosol. To counter this effect and maintain ROS production, neutrophils use the voltage-gated proton channel Hv1 (Hvcn1, VSOP), [11, 12], which regulates proton efflux from the cytosol [13, 14].

The Hv1 proton channel is a membrane protein that maintains cellular pH homeostasis by releasing protons across the plasma and the phagosome membranes. The protein has four transmembrane segments folded in a proton-conducting voltage-sensing domain (VSD) that resembles the corresponding domain of voltage-gated sodium, potassium, and calcium channels [15, 16]. While NOX2 activity is inhibited by membrane depolarization and cytosolic acidification, Hv1 is activated under these same conditions. In the absence of Hv1, neutrophil phagosomes were found to exhibit impaired ROS production [17] neutrophils from Hvcn1-/- mice also exhibit Ca^2+^ influx defects, resulting in impaired actin polymerization and neutrophil migration [18].

The importance of NOX2 in bacterial killing has been well described in patients with mutations in these subunits, most commonly gp91^phox^, who are highly susceptible to infection [19]. We also reported that mice with deletions in gp91^phox^ are unable to control blinding bacterial and fungal infections of the cornea [20, 21]. In the current study, we used *Hvcn1^−/−^*mice to examine the role of Hv1 in a clinically relevant murine model of blinding *Pseudomonas aeruginosa* corneal infection (keratitis). We found that corneal disease was more severe in infected *Hvcn1^−/−^* mice, which was associated with increased bacterial growth in the cornea and reduced neutrophil recruitment. However, contrary to our expectations, neutrophils from *Hvcn1^−/−^* mice produced more ROS than control neutrophils in response to *P. aeruginosa.* Collectively, these findings demonstrate that in response to bacterial infection, the role for Hv1/VSOP appears to be independent of ROS production.

## 2. MATERIALS AND METHODS

### 2.1 Mice

Male and female C57BL/6J mice aged 6-8 weeks were purchased from The Jackson Laboratory (Bar Harbor, ME). Two *Hvcn1^−/−^* breeding pairs were graciously provided from Dr. Long-Jun Wu and colleagues, Mayo Clinic, Rochester, MN [22]. Mice were housed in the University of California, Irvine vivarium. Age-matched, male and female mice were used for all experiments, and the experimental protocol was approved by the UC Irvine IACUC.

### 2.2 Bacterial strains and culture conditions

PAO1, PAO1-GFP and ΔpscD were obtained from Dr. Arne Rietsch (Case Western Reserve University). PAO1 is the parent strain that produces Type III secretion enzymes ExoS and ExoT, PAO1-GFP expresses the green fluorescent protein that enables us to visualize the bacteria in infected corneas and in neutrophils [23, 24], whereas Δ*pscD* does not produce the needle structure required for transfer of ExoS and ExoT to mammalian cells [25]. Bacteria were grown to mid-log phase (∼1× 10^8^ bacteria/ml) in high-salt Luria-Bertani (LB) broth, at 37°C with 5% CO_2_. Bacteria were washed and suspended in 1x PBS to a final concentration of 5×10^4^ bacteria/2 µl for *in vivo* infections and 3×10^8^ (MOI 30) for *in vitro* assays.

### 2.3 Murine model of *Pseudomonas aeruginosa* keratitis

We have been using a murine model of *P. aeruginosa* keratitis in which the corneal epithelium is abraded with 3 x 5mm parallel scratches using a sterile 30-gauge needle followed by topical application of a bacterial suspension in 2 µl PBS. While the inoculum depends on the strain *P. aeruginosa*, we use 5×10^4^ PAO1 [23, 24, 26, 27]. CFU were quantified after 2h to verify the inoculum for each experiment. After 24h and 48h, mice were euthanized by CO_2_ asphyxiation followed by cervical dislocation and were positioned in a 3-point stereotactic mouse restrainer for eye imaging. Corneal opacity (brightfield [BF]) and total bacteria (GFP) were visualized in the intact cornea using a high-resolution stereo fluorescence MZFLIII microscope (Leica Microsystems). ImageJ was then used to calculate corneal opacity and GFP intensity as we described in prior studies [23, 24]. Percent corneal opacity considers the area of opacity compared with the healthy transparent region. All images were obtained using the LAS V4.5 Software under the same magnification (×20). and corneal opacity and GFP were imaged and quantified by image analysis software (ImageJ, NIH).

### 2.4 Quantification of viable bacteria (colony forming units, CFU)

Whole eyes were collected and homogenized in 1 ml PBS using a 5mm steel ball bearing and Qiagen TissueLyser II at 30 Hz for 3 minutes. Serial log dilutions (10 µl) of the homogenate were plated on LB agar plates and incubated overnight at 37°C with 5% CO_2_. Colonies were counted manually, and viable bacteria were calculated as CFU/ml: number of colonies × dilution factor × 100.

### 2.5 Quantification of *in vivo* reactive oxygen species

Mice were infected with 5×10^4^ PAO1 in 2µl PBS (as detailed above). After 24h, mice were anesthetized, and a pocket was created in the stroma of the eye using a 30 g needle. A blunt needled Hamilton syringe was used to inject 2µl of dCFDA (2’,7’-dichlorofluorescein diacetate; Cayman Chemical) intrastromally. After 15 minutes, eyes were imaged for total ROS (GFP) and images were quantified by ImageJ (NIH).

### 2.6 Histology and immunohistochemistry (IHC)

Whole eyes were fixed in Davidson Fixative (Polysciences) for 24h and transferred to 70% ethanol. Paraffin embedding and sectioning was performed by the UCI Optical Microanatomy Core as part of the P30 Visual Sciences core facility grant. Sections (8 µM) were deparaffinized and hydrated before staining with hematoxylin and eosin (Fisher Scientific). Sections were imaged using Keyence BZ-X microscope with a Diagnostic Instruments camera attachment.

For immunohistochemistry, tissue sections were washed in PBS, permeabilized in 1% Tween-20 in PBS (PBST, Fisher Bioreagents), and incubated 1 hour with Fc block (BioLegend) and normal goat serum (Jackson ImmunoResearch) diluted in 1% BSA (Fisher Bioreagents). Rat Ly-6G/Ly-6C Monoclonal Antibody NIMP-R14 (Invitrogen) to detect neutrophils was diluted 1:50 in blocking buffer and incubated overnight at 4°C. Slides were washed 3x for 10 minutes with PBS. Goat anti-Rat 647 (Invitrogen) was diluted 1:1000 and added to sections for 1 hour at RT protected from light. Slides were then washed 4x for 10 minutes in PBS and counterstained with DAPI. Finally, slides were mounted using Vectashield® Antifade Mounting Medium (Vector Laboratories) and imaged on a Leica Stellaris SP8 confocal microscope.

### 2.7 Flow cytometry

Dissected corneas were incubated in 500 µl collagenase (3 mg/ml, C0130; Sigma Aldrich) in RPMI (Gibco), with 1% HEPES (Gibco), 1% penicillin-streptomycin (Gibco), 0.5% BSA (Fisher Bioreagents), and 2 µl of 1M Calcium Chloride for 1 h at 37°C. Recovered cells were incubated 10 min with anti-mouse CD16/32 Ab (BioLegend) to block Fc receptors. Cells were then incubated 20 min at 4°C with anti-mouse CD45-PE-Cy5, Ly6G-BV510, Ly6C-PE-Cy7, CD11b-PETxRed, CCR2-BV421, and F4/80-FITC (1:100 dilution, BioLegend) and fixable viability dye e780 (BD Biosciences). Cells were rinsed with FACS buffer and fixed with Cytofix/Cytoperm (BD Biosciences) for 15 min at 4°C, washed with PBS and resuspended in FACS buffer. Cells were then analyzed using the Novocyte Quanteon System 4 flow cytometer (Agilent), and data were analyzed using NovoExpress software.

### 2.8 Granule protein and cytokine quantification

Infected corneas were dissected, and underlying vascularized iris tissue was carefully removed and placed in a 2 ml tube containing 500 µl of sterile PBS (Corning) and a sterile 5mm steel ball bearing. Corneas were lysed using a Qiagen TissueLyser II at 30 Hz for 3 minutes. Samples were then centrifuged at 16,0000 rpm for 10 minutes at 4°C and granule proteins and cytokines were measured in cell-free supernatant. We used R&D Systems mouse DuoSet ELISA kits for IL-1β, CXCL1, CXCL2, myeloperoxidase (MPO) and neutrophil elastase according to the manufacturer’s protocol. ELISA plates were read on an Agilent (BioTek) Cytation5, and expressed as pg/ml cytokine. Cytokine production by stimulated bone marrow neutrophils were also assayed by ELISA The limit of detection of these antibodies is 1.5 pg/ml (CXCL1, CXCL2, myeloperoxidase), 4.8 pg/ml (IL-1β), and 2.27 pg/ml (neutrophil elastase).

### 2.9 Neutrophil Isolation

Mouse whole bone marrow was removed from femurs and tibias via centrifugation and was resuspended in 500 µl of chilled EasySep™ buffer (Stemcell). Bone marrow neutrophils were then isolated using the Stemcell EasySep™ Mouse Neutrophil Isolation Kit following the manufacturers protocol. Mouse neutrophils were counted manually and resuspended at 1×10^6^ in RPMI 1640 without phenol red (Gibco). This protocol routinely yields 80-90% purity.

### 2.10 HL60 cells

HL60 cells were originally derived from a patient with acute promyelocytic leukemia, and were purchased from ATCC (Gaithersburg, MD) and Synthego (Redwood City, CA). Cells were maintained in 10% FBS (Cytivia), RPMI 1640 with 25mM HEPES (Gibco). Cells were differentiated to a neutrophil phenotype for 5 or 6 days in 1.25% Dimethyl sulfoxide (DMSO) (Sigma Aldrich). Hv1 KO HL60 clones were generated by Synthego using proprietary Crispr-Cas9 gene editing technology. Details are provided in Supplemental Material.

### 2.11 Reactive Oxygen Species Assays

Murine bone marrow neutrophils and differentiated HL-60 cells were incubated at 37°C with 5% CO_2_ in RPMI 1640 without phenol red (Gibco) with 500 µM Luminol (Sigma Aldrich) for 30 minutes before stimulation (Luminol measures extra- and intracellular H_2_O_2_). Neutrophils were added to black sided, optically clear bottom 96-well plates (Corning) at 2×10^5^/well and DPI (Sigma Aldrich) was added to respective wells at a final concentration of 10 µM for 15 minutes prior to stimulation. *P. aeruginosa* (PAO1 or ΔpscD), PMA (100 ug/ml, Sigma Aldrich), and zymosan (100 ug/ml, Invivogen) were added and the plate was read on a Cytation5 plate reader (Agilent, BioTek) for 90 minutes at 37°C, taking luminescent (Lum) readings every 2 minutes. Time course curves were generated by the Cytation5 Gen5 software. ROS production in HL60 cells was measured under the same conditions as mouse neutrophils (on 96-well plate reader, 2×10^5^ cells/well), except the PMA concentration was 100 nM. The area under the curve was calculated from time courses using either GraphPad Prism or OriginPro software.

### 2.12 Neutrophil phagocytosis of *P. aeruginosa*

Neutrophils from the peritoneal cavity of C57BL/6 or *Hvcn1*-/- mice following sterile inflammation with casein (Sigma-Aldrich) were suspended at 4×10^5/well for 30min in RPMI with 2% FBS +/− cytochalasin D (Sigma-Aldrich). PAO1-GFP was added to the neutrophil suspension at MOI 30:1 for 15 mins after which gentamicin (400 µg/ml, Sigma-Aldrich) was added to kill extracellular bacteria. Infected neutrophils were then incubated 5 mins with Fc block in FACS buffer, and then with PE-Ly6G APC-CD11b to identify neutrophils (BioLegend), and eFluor780 viability stain (eBiosciences) for 30 mins and with DAP (Fisher). Cells were examined in the Amnis ImageStream, and the percent of neutrophils with associated (intracellular) GFP-bacteria was quantified using Amnis software.

### 2.13 Statistics

For *in vivo* experiments, statistical significance was determined using paired t-tests (GraphPad Prism). *In vitro* assays required a minimum of 3 biological repeats using the mean of four technical replicates. ROS production in HL60 cells stimulated with PMA required a minimum of four biological repeats using the mean of 2-3 technical replicates. Statistical significance was determined using unpaired t-tests (OriginPro). Error bars indicate mean ± SEM, unless otherwise indicated. p values less than 0.05 are considered significant. The number of biological replicates for each experiment is noted in the figure legends.

## 3. RESULTS

### Hv1 regulates bacterial killing and disease severity in a murine model of *Pseudomonas aeruginosa* corneal infection

Human and murine corneal infections are characterized by neutrophil infiltration, which together with bacterial growth, disrupt corneal transparency and lead to corneal opacification. This resolves only after the bacteria are killed by neutrophils (or by antibiotics). We reported that GP91^PHOX^ /CybB^−/−^ mice are unable to kill *P. aeruginosa* in infected corneas, resulting in more severe disease and demonstrating that NADPH oxidase and ROS are required to control bacterial growth [21]. As Hv1 sustains release of ROS generated by NOX2, we assessed the role of the Hv1 voltage-gated proton channel in *P. aeruginosa* keratitis. Corneas of C57BL/6 and *Hvcn1^−/−^* mice were abraded and infected topically with 5×10^4^ *P. aeruginosa* strain PAO1 that constitutively expresses green fluorescent protein (GFP-PAO1). After 24h or 48h, corneal opacity caused by infiltrating neutrophils and bacterial replication and GFP+ bacteria were quantified by image analysis, and CFU were counted as a measure of viable bacteria.

Although corneal opacification was observed in both strains of mice, infected *Hvcn1^−/−^* corneas had more severe corneal opacification (**Figure 1A,B)**. Total bacterial mass (GFP) and viable *P. aeruginosa* were higher in infected *Hvcn1^−/−^* corneas 24h and 48h post infection compared with C57BL/6 corneas (**Figure 1C)**.

**Figure 1.**
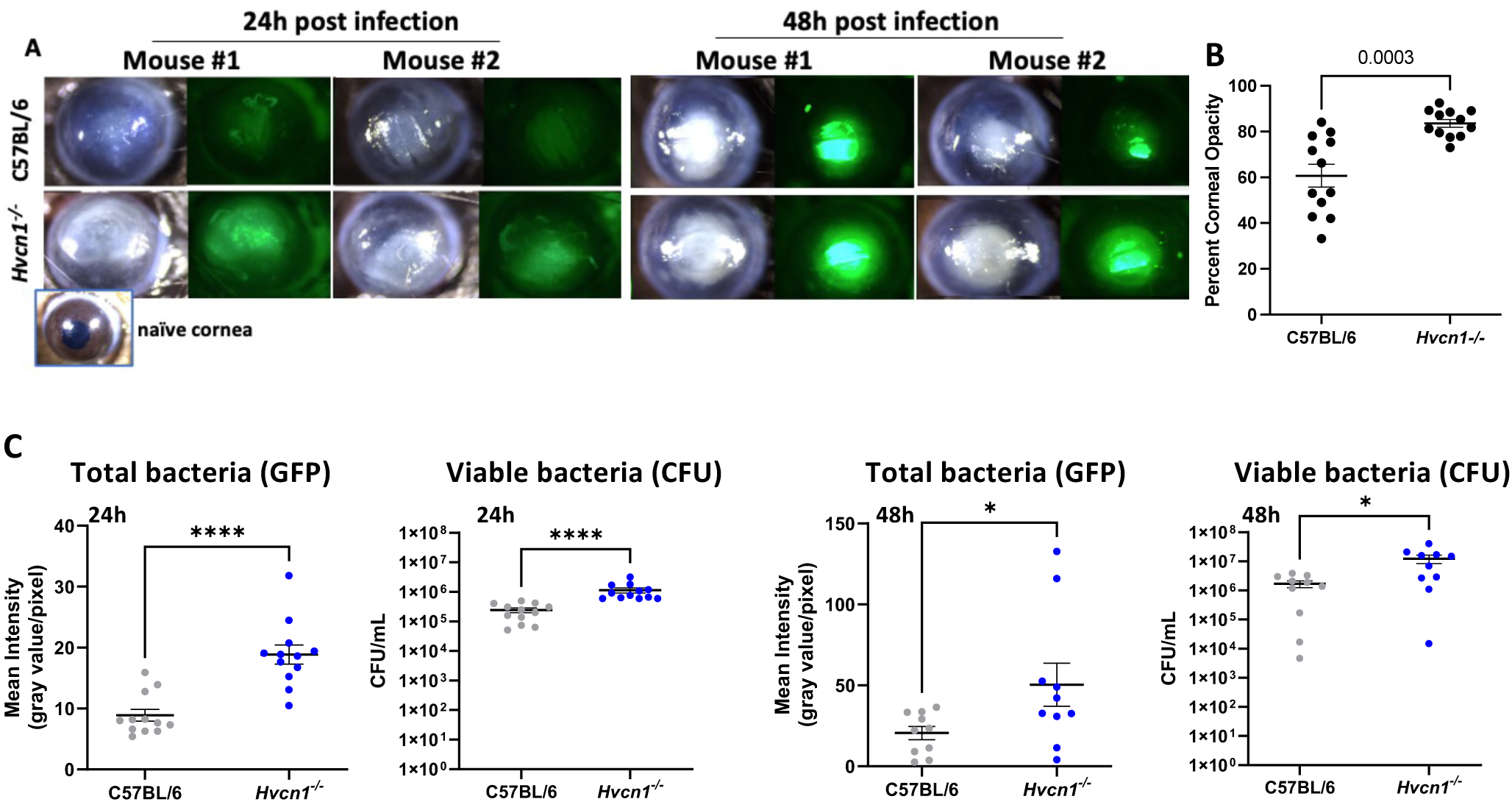
Role of Hv1 in *Pseudomonas aeruginosa* keratitis. Corneas of C57BL/6 and *Hvcn1^−/−^* mice 24h or 48h after topical infection with GFP-PAO1 bacteria. **A.** Representative images of corneal opacification (brightfield) and GFP-PAO1 in infected C57BL/6 and *Hvcn1^−/−^* mice. **Inset**: transparent cornea of a naïve mouse. **B.** Percent corneal opacity quantified by image analysis. **C.** Total GFP+ bacteria in infected corneas quantified by image analysis, and viable bacteria quantified by CFU 24h and 48h post infection. n=12, three experiments combined.

To determine if the impaired bacterial killing in *Hvcn1^−/−^* mice is associated with decreased ROS production, corneas of C57BL/6 and *Hvcn1^−/−^* mice were infected with PAO1 (not GFP expressing), and 2 µl dCFDA (2’,7’-dichlorofluorescein diacetate) was injected intrastromally after 24h as we described [20]. Total fluorescence was quantified by image analysis after 15 min.

There was significantly less fluorescence in infected *Hvcn1^−/−^* compared with C57BL/6 corneas (**Figure 2),** indicating less ROS. This is consistent with the role of ROS in killing *P. aeruginosa* in infected corneas.

**Figure 2.**
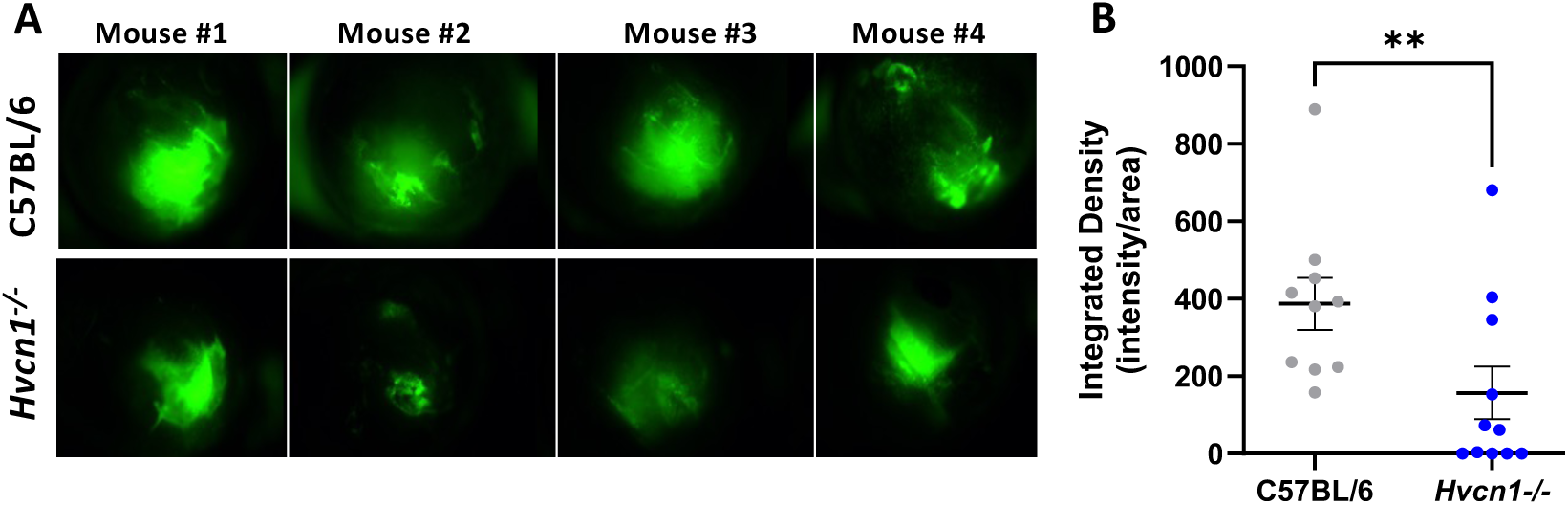
ROS production in infected corneas of C57BL/6 and *Hvcn1^−/−^* mice. 24h post infection with PAO1 (not GFP expressing), cDFDA was injected intrastromally and fluorescence imaged after 15 min. Representative corneas **(A)** and quantification of fluorescent cDFDA **(B).** Statistical significance was calculated using paired t-test, n=12, combined data from three independent experiments. Original magnification of images is x20.

### Impaired neutrophil recruitment to *P. aeruginosa* infected *Hvcn1^−/−^* corneas

The cornea is the anterior, transparent tissue that plays a critical role in allowing light to penetrate to the retina ((**Figure 3A).** Healthy, transparent human corneas have resident dendritic cells and macrophages, and recent studies also revealed the presence of T cells [28, 29]. Corneas of humans and mice are comprised of an external epithelial layer, the corneal stroma and the corneal endothelium that maintain transparency (**Figure 3B**); however, following *P. aeruginosa* infection, neutrophils rapidly infiltrate the stroma from limbal capillaries and from iris vessels resulting in increased corneal thickness as shown in histological sections and the presence of Ly6G+ neutrophils (**Figure 3C,D)**. While lower ROS production in infected *Hvcn1^−/−^*corneas may be due to the role of Hv1 in ROS production, decreased ROS may also be explained by number of neutrophils recruited to the site of infection. We therefore quantified the neutrophils in *P. aeruginosa* infected *Hvcn1^−/−^*corneas by flow cytometry.

**Figure 3.**
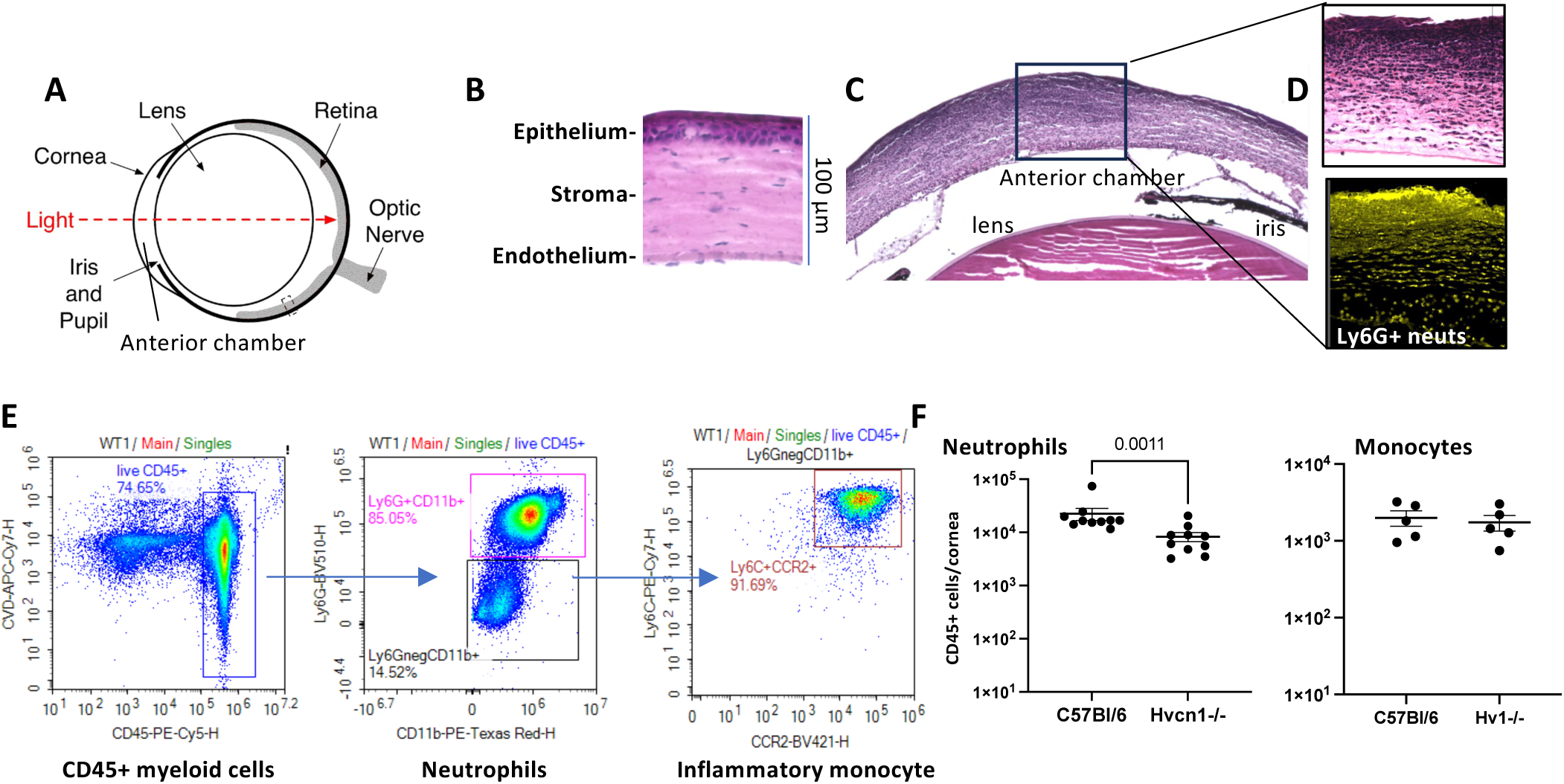
Neutrophil and monocyte recruitment to PAO1 infected corneas. **A**. Diagram of the mouse eye; **B.** Representative images of H&E - stained corneal sections of a naïve mouse (**B)** and infected C57BL/6 mouse corneas 24h post infection **(C,D). C.** Low power image of the central cornea of a 24h post infection with PAO1; **D.** Higher magnification of infected cornea by H&E (upper panel) to show total infiltrate and Ly6G+ neutrophils (lower panel). **E,F**: Infiltrating neutrophils and monocytes to infected corneas assessed by flow cytometry. **E.** Representative flow cytometry plots of live, single, CD45+ cells identifying neutrophils (Ly6G+CD11b+) and inflammatory monocytes (CD11b+, Ly6c+, CCR2+, Ly6G-). **F.** Quantification of total neutrophils and monocytes. n=6 infected mice, two experiments combined (data points represent individual infected corneas). All experiments were repeated x3 and combined or representative data are shown. Statistical significance was assessed using paired student’s t-test and p<0.05 was considered significant.

Corneas were dissected 24h post infection, collagenase digested, and cells were isolated and incubated with antibodies to detect neutrophils (CD45+/CD11b+/Ly6G+) and inflammatory monocytes (CD45+/CD11b+/CD11c+/CCR2+/Ly6G-). Gating on live, singlet cells we found that neutrophils comprised ∼85-90% of the total cellular infiltrate in infected C57BL/6 corneas, with 10-15% monocytes (**Figure 3E)**. While there was no significant difference in the percentage of infiltrating neutrophils or monocytes, there were significantly fewer total neutrophils in *Hvcn1^−/−^*compared with C57BL/6 corneas and no difference in the number of monocytes (**Figure 3F).**

These data indicate that Hv1 is required for recruitment of neutrophils to infected corneas, which explains the decreased ROS production shown in Figure 2 and indicates that Hv1 has a role in neutrophil migration to the cornea. This finding is consistent with the reported observation that *Hvcn1^−/−^* neutrophils exhibit an impaired response to chemotactic peptides [18].

### *P. aeruginosa* infected *Hvcn1^−/−^* corneas have less granule proteins compared with C57BL/6 corneas, but no difference in chemokines

*P. aeruginosa* infection of corneas induces pro-inflammatory and chemotactic cytokine production by resident macrophages, epithelial cells and keratocytes to initially recruit circulating neutrophils, which then produce these cytokines in response to replicating bacteria [27]. Activated neutrophils also release myeloperoxidase (MPO) and neutrophil elastase (NE) from primary granules. To assess the effect of Hv1 deficiency on MPO, NE and cytokine production in *P. aeruginosa* infected corneas, *Hvcn1^−/−^* and C57BL/6 corneas were infected with PAO1, and after 24h corneas were dissected and subjected to bead homogenization. Following centrifugation, granule proteins and cytokines were quantified by ELISA.

As shown in **Figure 4**, there was significantly less MPO and NE in infected *Hvcn1^−/−^* compared with C57BL/6 corneas. While this likely reflects the lower number of neutrophils in *Hvcn1^−/−^* corneas, production of neutrophil chemokines CXCL1 and CXCL2 and the pro-inflammatory cytokine IL-1β was not significantly different. Neutrophils are the only source of MPO and NE, but cytokines are produced by other cells in the cornea, including resident macrophages, epithelial cells and keratocytes, and infiltrating monocytes, The apparent selective role for Hv1 on secretion of granule proteins is also consistent with reports that *Hvcn1^−/−^* neutrophils produce elevated MPO and NE compared with WT neutrophils in response to stimulation with phorbol myristic acid (PMA) [30].

**Figure 4.**
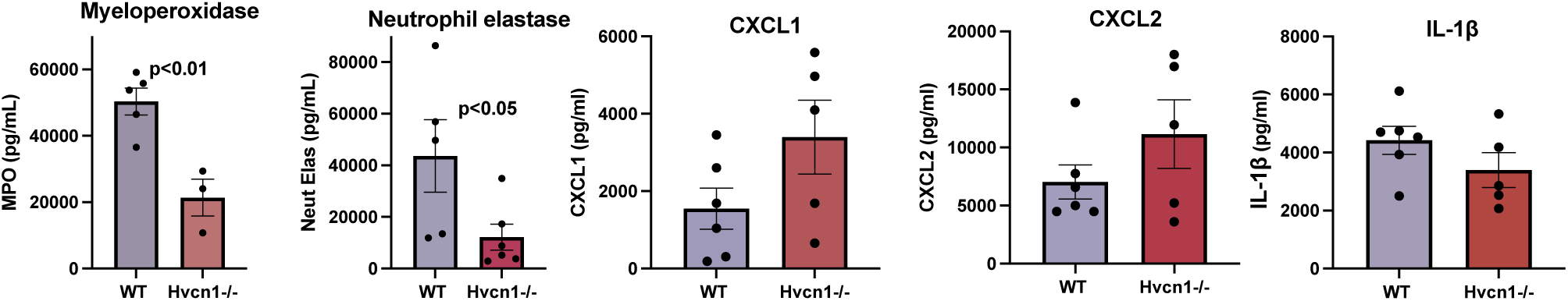
Granule protein and cytokine production in *Hvcn1^−/−^* infected corneas. Corneas of C57BL/6 and *Hvcn1^−/−^* mice were infected with *P. aeruginosa* and after 24h, corneas were homogenized and neutrophil granule proteins myeloperoxidase (MPO) and neutrophil elastase, pro-inflammatory cytokines and chemokines were quantified by ELISA. Data are representative of 3 repeat experiments. Statistical significance was calculated using paired t-test, and p<0.05 was considered significant.

### *Hvcn1^−/−^* neutrophils corneas produce elevated ROS in response to *P. aeruginosa*

We reported that *P. aeruginosa* Exoenzyme S inhibits NOX2 activity and ROS production by human neutrophils by ADP ribosylation of Ras, thereby blocking recruitment of NOX2 cytosolic proteins to the membrane and inhibiting NOX2 assembly [21]. Consequently, infection with the T3SS injectosome mutant Δ*pscD,* which does not form a needle structure and therefore cannot inject ExoS or other exoenzymes into the host cell cytosol, cannot inhibit NOX2 assembly and ROS production by primary human neutrophils [21].

To examine the role of Hv1 on *P. aeruginosa* induced ROS production, we infected neutrophils from *Hvcn1^−/−^* and C57BL/6 mice with the parent PAO1 strain, the Δ*pscD* mutant or with phorbol myristate acetate (PMA) as a known ROS inducer. In some assays, NOX2 inhibitor Diphenyleneiodonium (DPI) was included to inhibit ROS production. Neutrophils were then incubated for 90 min in the presence of Luminol.

As expected from our earlier studies, C57BL/6 neutrophils infected with Δ*pscD*, but not the parent PAO1 strain, induced extracellular ROS production **(Figure 5A)**. Given the reported role of Hv1 in promoting ROS production, we anticipated that ROS production would be lower in the absence of Hv1. However, we instead found significantly elevated ROS production by *Hvcn1*^−/−^ compared with C57BL/6 neutrophils infected with Δ*pscD* or stimulated with PMA **(Figure 5B).** Quantification of the area under the curve in repeat experiments showed consistently elevated ROS production in *Hvcn1*^−/−^ compared with C57BL/6 neutrophils **(Figure 5C).** *Hvcn1*^−/−^ neutrophils also produced elevated ROS in response to stimulation with zymosan (**Figure S1A).** The increased ROS was not due to differences in cell numbers as neutrophils from both mouse strains produced the same amount of IL-1β following LPS/ATP or LPS/PAO1 stimulation (**Figure S1B).**

**Figure 5.**
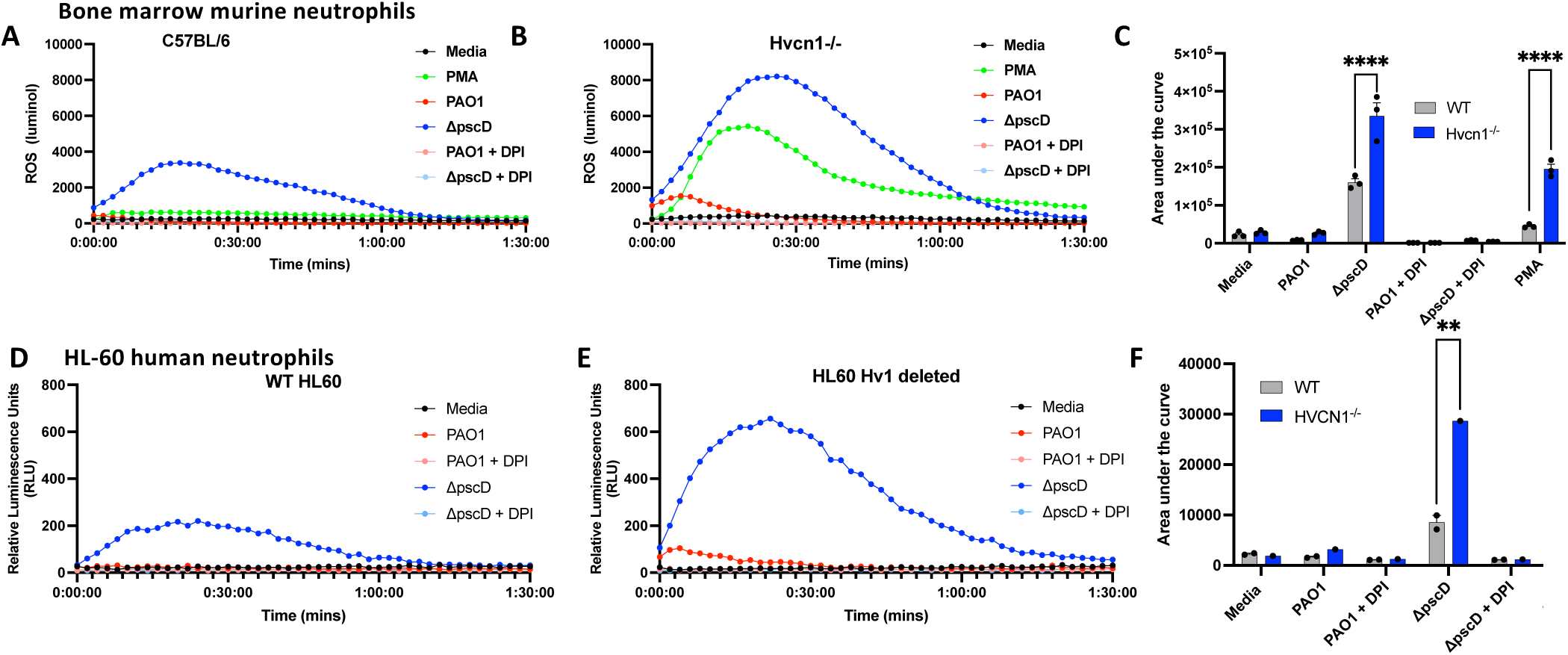
Reactive oxygen production by *Hvcn1^−/−^* neutrophils. **A-C**: Murine bone marrow neutrophils from C57BL/6 and *Hvcn1^−/−^* mice were incubated with 500 µM luminol and stimulated with parent strain *P. aeruginosa* PAO1, the type III secretion mutant Δ*pscD* or with 100 µM phorbol myristate acetate (PMA) +/− DPI (**C,D**). **A, B.** representative time course of ROS production by C57BL/6 and *Hvcn1^−/−^* neutrophils measured as relative fluorescent units (RFU) per second; **C.** Total ROS production defined as area under the curve. **D-F.** Parent and Hv1 CRISPR deleted HL-60 human myelocyte cell line following differentiation to neutrophils. **D, E**. Representative time course of ROS production; **F.** Area under the curve calculated from 3-6 biological replicates. Total ROS production was quantified as area under the curve. Statistical significance was calculated by ANOVA with Tukey’s post hoc analysis.

**Figure 6.**
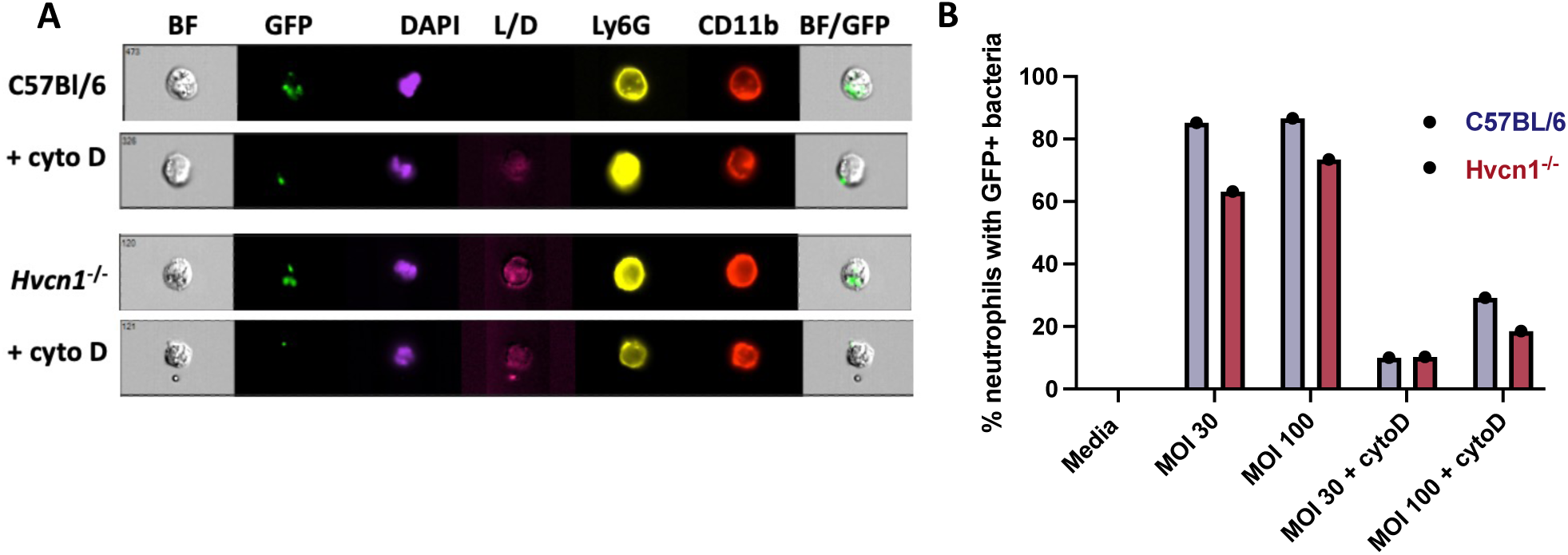
Phagocytosis of *P. aeruginosa* by *Hvcn1^−/−^* neutrophils. **A-C**: Murine peritoneal neutrophils from C57BL/6 or *Hvcn1^−/−^* mice were incubated 15 min with GFP-PAO1 (30:1 or 100:1 MOI) in the presence or absence of cytochalasin D (CytoD). Gentamycin was added to kill extracellular bacteria, and neutrophils were stained with antibodies to Ly6G (PE conjugated), APC CD11b, and the e780 live/dead stain and DAPI. Cells were then imaged and quantified by Amnis Imagestream. **A.** Representative images; **B.** quantification of percent neutrophils with GFP-PAO1. Experiments were repeated twice.

As an alternate approach, the Hvcn1 gene in the HL60 human myeloid cell line was mutated using CRISPR-Cas9 and differentiated into neutrophil-like cells following 5 days incubation with DMSO. Parent and Hv1-deleted cells were then incubated with *P. aeruginosa* strains PAO1 or Δ*pscD* or with PMA in the presence of luminol. As with murine *Hvcn1*^−/−^ neutrophils, ROS production was elevated in Hv1-deleted cells compared with parent HL60 cells (**Figure 5C-E**).

Supplemental **Figure S2**. shows that PMA induces ROS in differentiated but not undifferentiated HL60 cells, and that Hv1 deletion has no effect on PMA-induced ROS. Collectively, these findings indicate that under these conditions, Hv1 has a suppressive role on ROS production by *P. aeruginosa* and other stimuli.

### *Hvcn1^−/−^* neutrophils have no defect in phagocytosis of *P. aeruginosa*

To determine if impaired bacterial killing in infected corneas is consequence of less phagocytosis in the absence of Hv1, peritoneal neutrophils were isolated from C57Bl/6 and *Hvcn1*^−/−^ mice and incubated 15 minutes with GFP-PAO1 at a multiplicity of infection (MOI) of 30:1 or 100:1. Neutrophils were then washed with PBS containing gentamycin to kill extracellular bacteria, and the number of neutrophils with bacterial were detected using Imaging flow cytometry, Neutrophils were identified as Ly6G+, CD11b+ cells. As a negative control for phagocytosis, cells were also incubated in the presence of the actin polymerization inhibitor cytochalasin D.

GFP-PAO1 were detected in C57Bl/6 and *Hvcn1*^−/−^ neutrophils, and cytochalasin D inhibited bacterial uptake (**Figure 5A)**. However, there was no significant difference between C57Bl/6 and *Hvcn1*^−/−^ neutrophils at either MOI (**Figure 5B)**. Therefore, the observed reduction in bacterial killing in *Hvcn1-/-* infected corneas is not due to impaired phagocytosis by neutrophils.

## DISCUSSION

Hv1 is the only known voltage-gated proton channel in mammals. In contrast to other voltage-gated channels, it does not possess a pore domain [11, 12], and Hv1 is also unique in how it responds to mechanical stress [31]. We reported that NADPH oxidase and ROS production is critical for controlling *P. aeruginosa* growth by neutrophils *in vitro* and in a clinically relevant murine model of blinding corneal infection where neutrophils comprise >80% of the total cellular infiltrate [7, 24]. In the current study, we anticipated that given the known role for Hv1 in facilitating ROS production, Hv1 deficiency would result in impaired ROS production and resulting failure to control bacterial growth in *P. aeruginosa* infected corneas. However, while we found impaired bacterial killing and more severe corneal disease in infected *Hvcn1*^−/−^ mice, the underlying reason was a defect in neutrophil recruitment to infected corneas as shown by immunohistochemistry and flow cytometry. These results were unexpected and underscore the importance of Hv1 in neutrophil trafficking rather than ROS production.

Our findings are consistent with those of El Chemaly *et a*l, and Okochi *et al* who showed *a* requirement for Hv1 in migration of neutrophils in response to agonists of formyl peptide receptors (N-formyl methionyl-leucyl-phenylalanine, fMLF) [18] [30]. Our *in vivo* data are consistent with those of Ramsey et al who showed data showing significantly more *Staphylococcus aureus* CFU in *Hvcn1^−/−^* mice 6h after intraperitoneal infection with 1 × 10^9^ (though not 1 × 10^8^) bacteria [12]. Zhao and Goldstein also showed that inhibiting Hv1 using a small C6 peptide resulted in significantly less recruitment of neutrophils to the lungs in a murine model of LPS – induced pulmonary inflammation [32, 33]. Together, these studies indicate that Hv1 promotes neutrophil migration in murine models of infection and inflammation. Our findings also extend the role of Hv1 to neutrophil trafficking in corneal infections.

While these studies show impaired recruitment of cells in *Hvcn1*^−/−^ mice, Okochi and colleagues reported that *Hvcn1*^−/−^ mice infected intranasally with *Candida albicans* had increased rather than decreased cellular infiltration to the lungs, although there was no effect on fungal killing [30]. Similarly, using a murine model of ovalbumin induced lung allergy, where ovalbumin is instilled into the lungs after prior sensitization, Du et al reported increased inflammation and cytokine production in *Hvcn1*^−/−^ mice [34].

Hv1 deficient mice were independently generated by Okochi et al [17] and by the Chapham group in the USA [14], which were used in the current study. Both used the same gene-trap strategy to disrupt the Hv1 gene, so the different findings are unlikely to be related to the mouse strains used. Instead, the different outcomes may relate to acute disease and infection models compared with longer term models. The current study, and those of Ramsey and of Zhao, examined responses in hours or days [14, 32]. In contrast, the other studies examined *C. albicans* lung infection or ovalbumin induced pulmonary inflammation at later time points when adaptive immunity is likely also involved [30, 34].

Based on prior reports that Hv1 facilitates NOX2 activity [14, 18], we anticipated that ROS production by *Hvcn1*^−/−^ neutrophils would be impaired; instead we found increased ROS production in response to stimulation with *P. aeruginosa*, zymosan or PMA. Similarly, following CRISPR-Cas9 deletion of Hv1 in the HL60 neutrophil cell line, we found increased ROS production compared with Hv1 expressing cells stimulated with *P. aeruginosa*. Our findings are in agreement with Okochi *et a*l, who reported increased rather than decreased ROS production by *Hvcn1*^−/−^ neutrophils in response to fMLF [35]. In addition, MDA-MB-231 breast cancer cells with deleted Hv1 produce more ROS than corresponding Hv1-expressing control cells [36], further indicating that ROS modulation by Hv1 depends on which cell types and stimuli are being examined.

The rationale for investigating the role of Hv1 in *P. aeruginosa* corneal infection was based on our earlier report showing that NOX2 activation and ROS production by neutrophil plays an essential role in killing bacteria in the cornea [21]. In that study, we showed that in the absence of GP91^PHOX^, neutrophil infiltration was increased, but neutrophils were unable to kill the bacteria. We reported that the *P. aeruginosa* Type III secretion system is essential for virulence in this model [26]; however, in the absence of a functional NOX2, infection with Type III secretion system mutants (including Δ*pscD*) were able to replicate in the cornea and cause disease [21], thereby demonstrating the importance of ROS production in controlling bacterial growth in the cornea. The resistance of T3SS expressing *P. aeruginosa* to neutrophils killing is based on the ADP ribosyl transferase (ADPRT) activity of Type III secretion proteins ExoS and ExoT, which inhibit ROS production by neutrophils. ExoS ADPRT specifically targets Ras that is required for recruitment of cytosolic PHOX proteins to the membrane associated proteins [21, 26].

We anticipated that corneal infection of *Hvcn1*^−/−^ mice would demonstrate a similar phenotype; however, whereas corneal infection of *Hvcn1*^−/−^ mice also resulted in increased bacterial growth, this was a consequence of impaired neutrophil recruitment to the cornea compared with C57BL/6 and GP91^PHOX-/-^ mice. These findings are consistent with reports showing that *Hvcn1*^−/−^ neutrophils exhibit an impaired ability to migrate in response to chemotactic peptides [18].

Taken together, our study adds to a growing body of evidence that the role of Hv1 in neutrophil-mediated host defense is highly context-dependent, influenced by the type of pathogen, site of infection, and the immune-modulatory mechanisms at play. In *P. aeruginosa* keratitis, Hv1 plays a dual role in supporting antimicrobial defense and regulating neutrophil recruitment, with the latter being more critical in our model. These findings not only expand our understanding of Hv1’s function in ocular immunity but also support further investigation of Hv1 inhibitors as adjunctive therapies. By modulating neutrophil activity without broadly suppressing immune function, Hv1-targeted approaches may offer a more defined strategy for managing infectious inflammation while minimizing tissue damage and preserving vision.

## Author contributions

PR, SA, JA, JTC, CZ and MEM performed all experiments and generated graphs and images. PR, FT and EP designed the experiments and wrote the manuscript.

## Acknowledgements

We would like to thank Dr. Liang Hong (University of Illinois, Chicago) for providing *Hvcn1^−/−^* breeding pairs. We would also like to thank Dr. Arne Rietsch at Case Western Reserve University for providing the *P. aeruginosa* strains used in this study.

---

**Figure S1.**
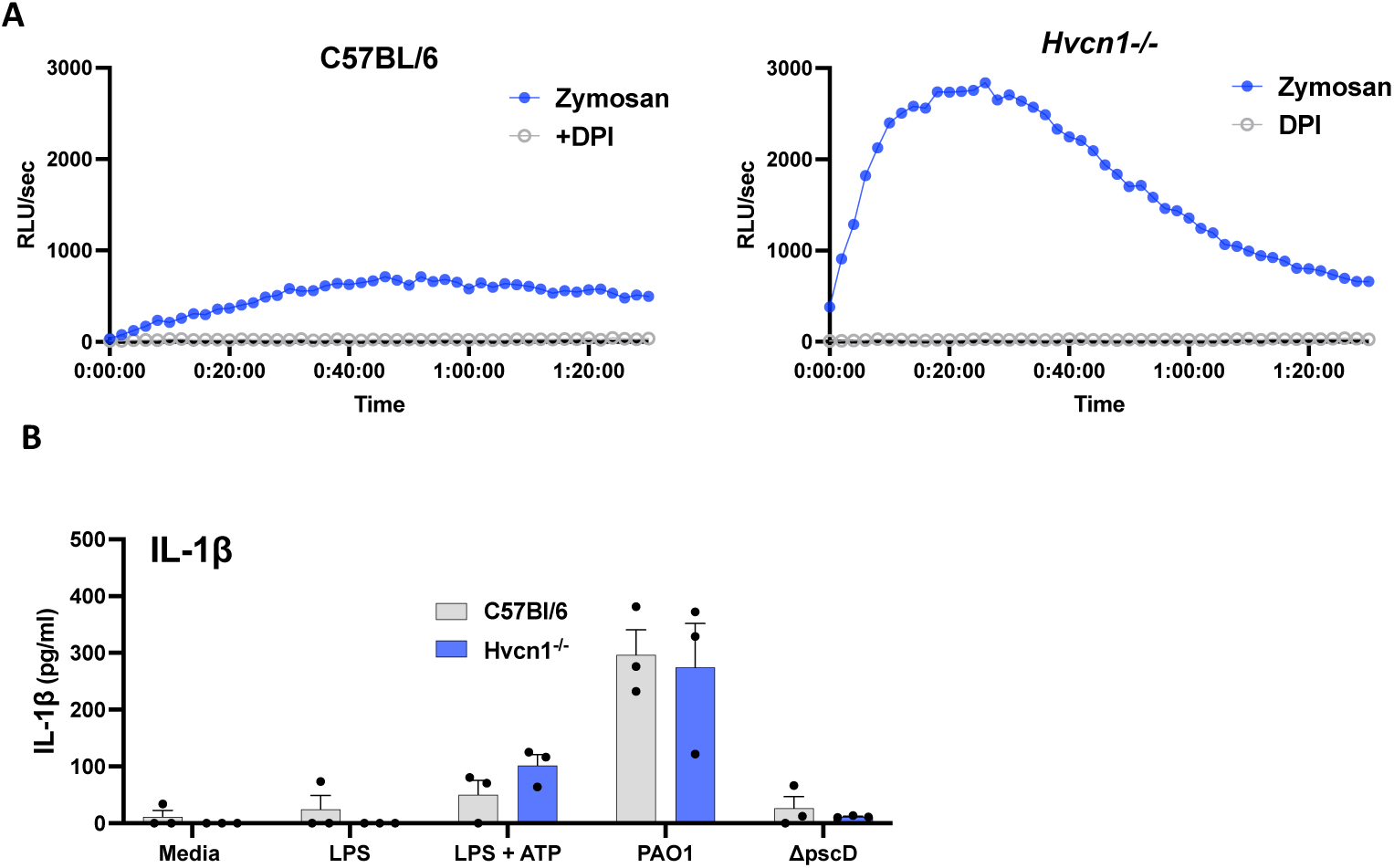
**A.** ROS production following incubation with zymosan +/− DPI. **B.** IL-1β secretion by C57BL/6 and *Hvcn1^−/−^* neutrophils incubated 3h LPS followed by 1h incubation with ATP, PAO1 or Δ*pscD* to activate the NLRP3 inflammasome.

**Figure S2.**
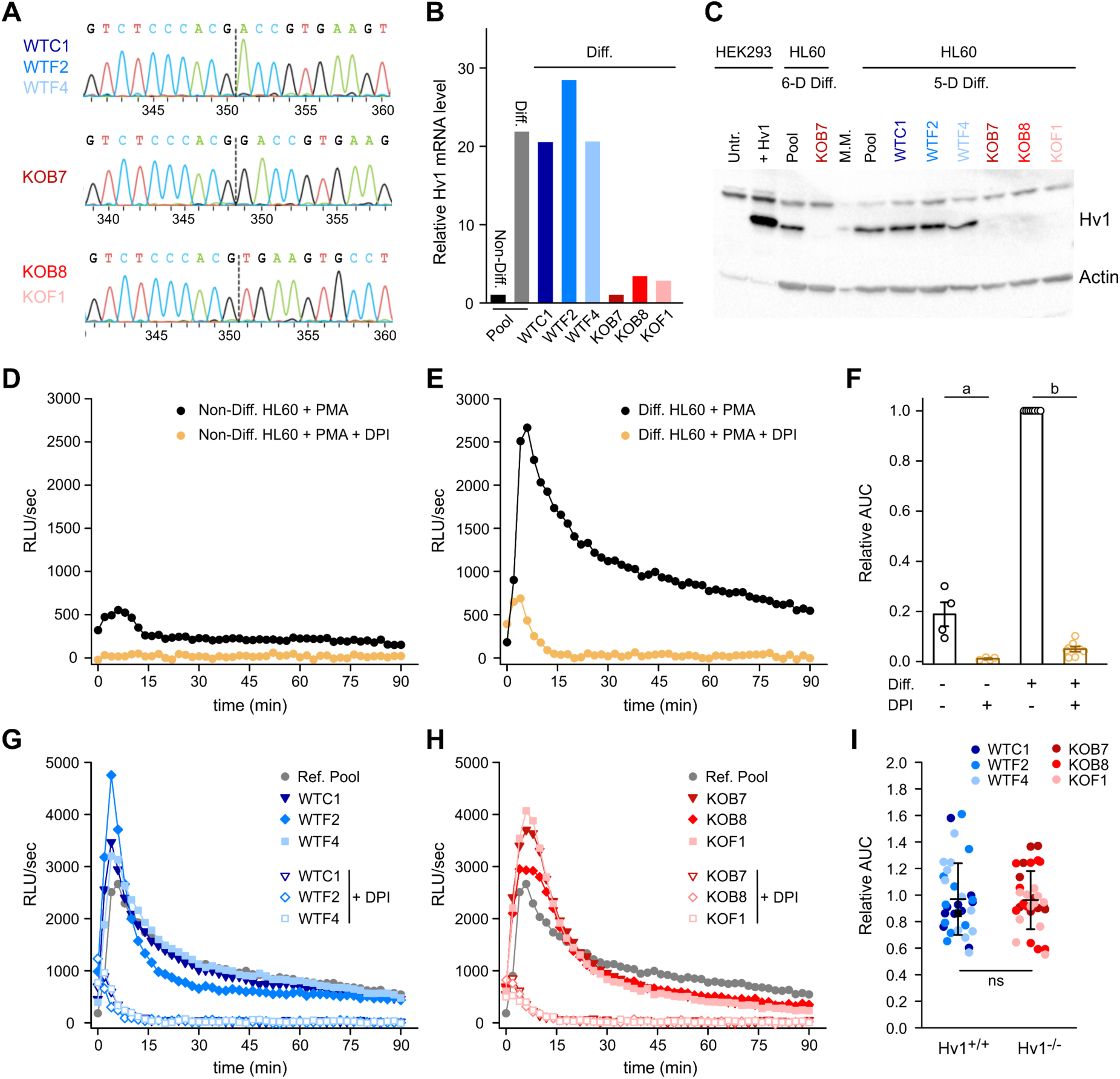
ROS production by Hv1 deleted HL60 cells in response to PMA stimulation. **A.** Sequence of the HVCN1 gene in the region of Crisper/Cas9 gene editing. (Top) Sequence shows clones that did not result in a change from wild type. (Middle) Sequence shows knock out (KO) clone B7 with an indel of a one base pair insertion and (Bottom) clones B8, F1 with the same indel of a 4 bp deletion. **B.** HVCN1 mRNA levels in edited and non-edited HL60 cells differentiated to neutrophil-like cells with DMSO treatment (1.25%) for 5 days, compared to undifferentiated and unedited HL60. Quantified by qPCR. **C.** Western blot with Hv1 and actin antibody shows expression in HL60 cells differentiated for 5 or 6 days with loss of expression in KO clones. HEK293 cells, untransfected (Untr.) or transfected with an Hv1 overexpression plasmid confirms antibody recognition of Hv1 protein band. **D-F.** ROS production in undifferentiated (**D**) and differentiated HL60 cells (**E**), after stimulation with 100 nM PMA. Cells were differentiated for 5 days and ROS were measured in the absence or presence of 10 µM DPI. **F.** Quantification of ROS production measured as relative area under the curve (AUC) of time courses like those shown in (**D, E**). Error bars are ±SEM. Non-Diff: n=4, p=0.0039 (a). Diff: n=8, p=9.7×10^−4^ (b). Data points represent repeat experiments. **G-I.** ROS production in WT (**G**) and Hv1 KO HL60 clones (**H**) measured under the same conditions as in (**D, E**). **I.** Quantification of ROS production as AUC of time courses like those shown in (**G, H**). Error bars are ±SD. WT: n=32, KO: n=33, (p=0.89, ns). Data points represent repeat experiments.

## Supplemental methods

### WT and Hv1 KO HL60 clones

Independent WT and Hv1 KO clones were isolated by single cell dilution from a pool generated from parental HL60 electroporated with chemically modified target-specific sgRNA and purified SpCas9 nuclease in the form of ribonucleoprotein complex and without selection markers. The sgRNA sequence was UUAAGGCACUUCACGGUCGU (cut location chr12:110,661,381). Editing was confirmed by Sanger sequencing (F-primer: GCATTCCCAGCACGGTACA, R-primer: CCGCCTTCTAGGCAGTCAC). An additional wild-type cell pool was generated from parental cells electroporated with SpCas9 only and confirmed to be unedited at target locus.

### Evaluation of Hv1 expression in HL60 cells

Hv1 mRNA levels were assessed using TaqMan Gene Expression Assays with human Hvcn1, Hs00260697_m1 and normalized with human GAPDH Hs00266705_g1. (ThermoFisher). For Western blots, HL60 and HEK293 cells were washed twice in ice-cold PBS with 1 mM PMSF (Millipore), centrifuging at 150 g for 5 min at 4C. The pellets were lysed in RIPA buffer (Thermo Scientific), containing 1 mM EDTA, 1 mM AEBSF (TCI America), 5 mM PMSF (Millipore), 1x HALT protease inhibitor (ThermoFisher) for 30 min on a rotator at 4C, centrifuged at 13,200 g × 10 min at 4C.

Soluble lysates were removed and protein concentration was determined using the Pierce BCA assay (Thermo Sci.). Lysates were loaded on Novex Wedgwell 4-20% gradient, Tris-Glycine gels (ThermoFisher) and electrophoresis was carried out at 150 V x 75 min. Proteins were wet-transferred at 100 V x 1.5 hours in Transfer Buffer (25 mM Tris, 192 mM Glycine, 20% Methanol) onto a PVDF Transfer Membranes (Thermo Sci.). Blots were blocked in 5% Nonfat Dry Milk (NFM) (Carnation) in 20mM Tris pH 7.5, 150mM NaCl, 50mM KCl, 0.1% TWEEN-20 (TBST) at RT for 15 min and probed with anti-Hv1 antibody (Invitrogen/ThermoFisher PA5-21008, 1mg/ml) at 1:1,000 overnight at 4C, washed 3x TBST and detected with goat anti-rabbit-HRP (Thermo Sci. 31480, 1 mg/ml) at 1:10,000 for 1.5 hour at RT. After imaging for Hv1, blots were treated with Restore PLUS Western Blot Stripping Buffer (Thermo Sci.) for 10 min, washed twice with water, blocked for 15 min in 5% NFM in TBST and re-probed with anti-β-Actin antibody-HRP (ThermoFisher MA515739HRP,1 mg/ml) at 1:100,000 for 1 hour at RT. After washing 3 times in TBST, bands were detected with Thermo SuperSignal West Femto Max sensitivity, using a Molecular Imager (Bio-Rad Chemi-Doc XRS System) per manufacturer’s directions.

